# Pathogenic TDP-43 Disrupts Axon Initial Segment Structure and Neuronal Excitability in a Human iPSC Model of ALS

**DOI:** 10.1101/2022.05.16.492186

**Authors:** Peter Harley, Guilherme Neves, Federica Riccio, Carolina Barcellos Machado, Aimee Cheesbrough, Lea R’Bibo, Juan Burrone, Ivo Lieberam

**Author notes:** Senior authors (Juan Burrone, Ivo Lieberam).

## Abstract

Dysregulated neuronal excitability is a hallmark of amyotrophic lateral sclerosis (ALS). We sought to investigate how functional changes to the axon initial segment (AIS), the site of action potential generation, could impact neuronal excitability in a human iPSC model of ALS. We found that early (6-week) ALS-related TDP-43^G298S^ motor neurons showed an increase in the length of the AIS, relative to CRISPR-corrected controls. This was linked to neuronal hyperexcitability and increased spontaneous contractions of hiPSC-myofibers in compartmentalised neuromuscular co-cultures. In contrast late (10-week) TDP-43^G298S^ motor neurons showed reduced AIS length and hypoexcitability. At a molecular level aberrant expression of the AIS master scaffolding protein Ankyrin-G, and the AIS-specific voltage-gated ion channels SCN1A (Nav1.1) and SCN8A (Nav1.6) mirrored these dynamic changes in excitability. Finally, at all stages, TDP-43^G298S^ motor neurons showed compromised activity-dependent plasticity of the AIS, further contributing to abnormal excitability. Our results point toward the AIS as an important subcellular target driving changes to neuronal excitability in ALS.

## Introduction

Amyotrophic lateral sclerosis (ALS) is a fatal neuromuscular disease, characterised by progressive degeneration of motor neurons (MNs) in the brain and spinal cord (Hardiman et al., 2017). In the majority of patients with ALS, TAR DNA-binding protein (TDP-43) mis-localises to the cytoplasm and forms ubiquitinated and hyperphosphorylated aggregates (Neumann et al., 2006; Sreedharan et al., 2008), which has been linked to abnormal RNA metabolism and stability, as well as changes to RNA splicing and gene expression (Hardiman *et al*., 2017; Narayanan et al., 2013). Dysregulated neuronal excitability is another key pathological hallmark of ALS (Gunes et al., 2020). Peripheral axonal and motor unit hyperexcitability, associated with spontaneous muscle fasciculations, are often observed in the early stages of the disease and have been shown to correlate with increased disease severity and reduced survival time (Kanai et al., 2006; Piotrkiewicz et al., 2008; Shimizu et al., 2014; Vucic and Kiernan, 2006). Conversely, neuronal hypoexcitability has been reported at later stages of the disease (Devlin et al., 2015; Gunes *et al*., 2020; Martinez-Silva et al., 2018). However, it remains unclear how pathogenic TDP-43 drives these changes to neuronal excitability.

The axon initial segment (AIS) is a specialised region of the proximal axon where action potentials are initiated (Kole et al., 2008). This subcellular domain is characterised by a high density of voltage gated sodium and potassium channels, particularly Nav1.1 (SCN1A), Nav1.2 (SCN2A), Nav1.6 (SCN8A) and KCNA1 (Kv1.1) that are anchored to the membrane by a unique cytoskeletal arrangement of scaffolding proteins including BIV-Spectrin and the master organiser Ankyrin-G (ANK3) (Hu et al., 2009; Leterrier, 2018). ANK3 can be alternatively spliced into an AIS-specific 480kDa isoform or shorter non-AIS 270kDa and 190kDa isoforms that play critical roles in myelination and dendritic spine morphology respectively (Nelson and Jenkins, 2016). Previous work has shown that both the length and position of the AIS can be modulated in an activity-dependent manner to fine-tune neuronal excitability (Evans et al., 2015; Grubb and Burrone, 2010; Jamann et al., 2021; Kuba et al., 2010; Sohn et al., 2019; Wefelmeyer et al., 2015). Modulation of AIS structure has been proposed as a homeostatic form of plasticity that acts to stabilise neuronal output and prevent abnormal levels of network activity. In this context, the AIS stands out as an attractive subcellular target for driving abnormal neuronal excitability in ALS.

In this study we sought to determine whether pathogenic TDP-43 could alter neuronal excitability by disrupting the AIS at a molecular, structural and functional level. We differentiated ALS patient-derived TDP-43^G298S^ hiPSCs into MNs and employed CRISPR-Cas9 to generate gene-corrected isogenic controls. We found that in early (6-week) MNs, pathogenic TDP-43^G298S^ triggered an increase in the expression of ANK3 (Ankyrin-G), SCN1A (Nav1.1) and SCN8A (Nav1.6) as well as increasing in the ratio of the AIS-specific 480kDa ANK3 isoform relative to shorter non-AIS isoforms. These molecular changes were linked to increased AIS length, neuronal hyperexcitability and increased spontaneous myofiber contractions in compartmentalised microfluidic neuromuscular co-cultures. In contrast, in late (10-week) MNs, pathogenic TDP-43^G298S^ caused TDP-43 mis-localisation, and reduced expression of ANK3, SCN1A and SCN8A, as well as an increase in the ratio of the non-AIS 190kDa ANK3 isoform over the AIS-specific 480kDa isoform. These changes were linked to reduced AIS length and neuronal hypoexcitability. Finally, at all stages, activity dependent plasticity of the AIS was compromised, further destabilising neuronal excitability. Taken together our findings point toward a unique and dynamic role of the AIS in modulating changes to neuronal excitability in ALS.

## Results

### 1. Pathogenic TDP-43^G298S^ causes increased AIS length and neuronal hyperexcitability in early MNs

We set out to determine whether pathogenic TDP-43^G298S^ could alter MN excitability by disrupting AIS structure and plasticity. We generated synchronised cultures of hiPSC-derived MNs by using Magnetic Activated Cell Sorting (MACS) to purify post-mitotic HB9+ MNs and matured them in defined co-cultures with GDNF-expressing astrocytes (Machado et al., 2019) (Figure 1b, Supplementary figure 1d). We employed CRISPR-Cas9 to generate TDP-43^G298S^ corrected hiPSCs (Fig. 1b, Supplementary Figure 1a). We then reconstructed the AIS of early 6-week MNs based on immunofluorescence staining of Ankyrin-G (Fig. 1c). We found that TDP-43^G298S^ triggered an increase in the length of the AIS relative to corrected and wildtype MNs (Fig. c,d). AIS diameter and start position from the soma remained constant (Supplementary figure. 2a,b).

**Figure 1.**
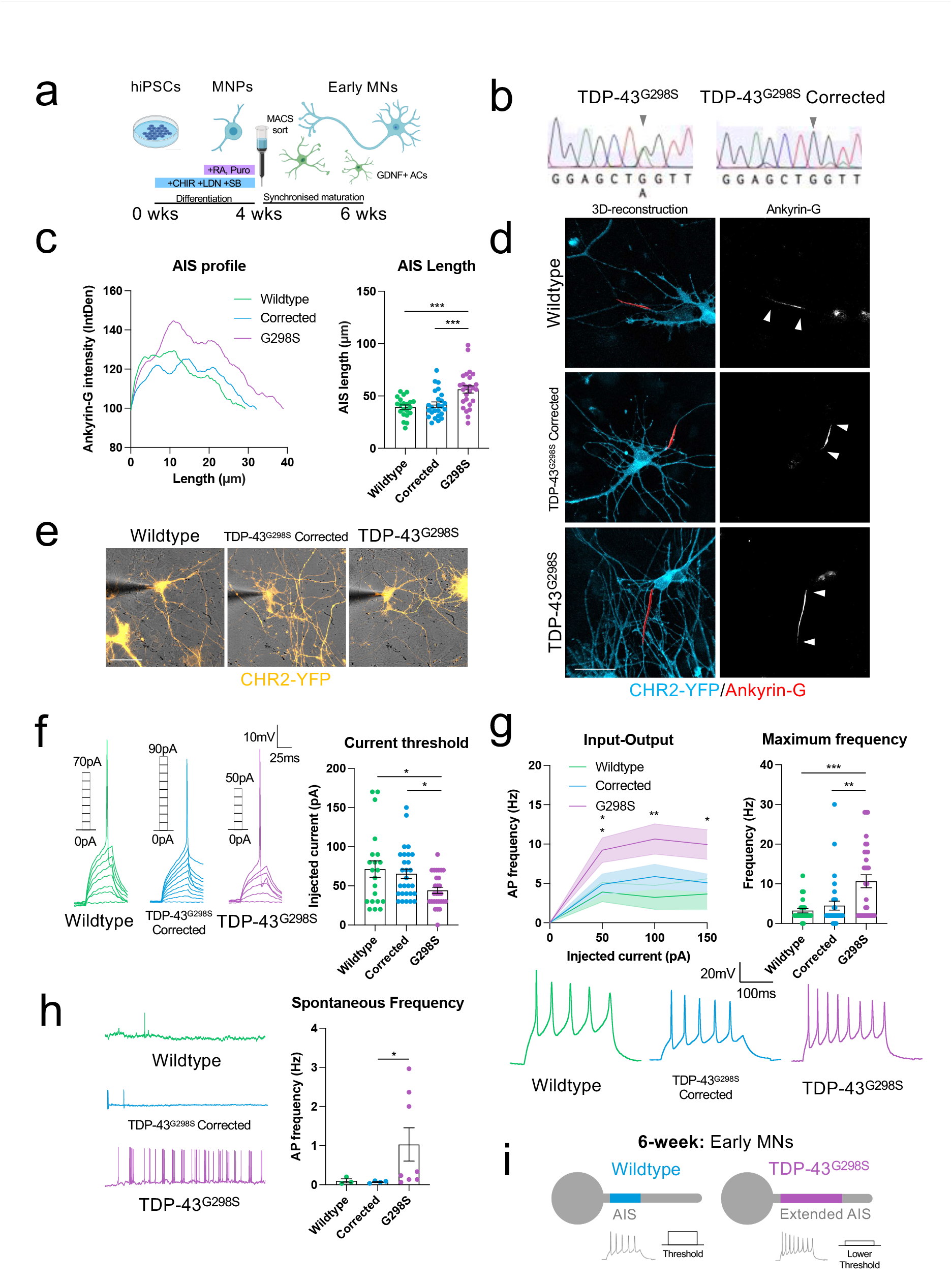
Pathogenic TDP-43^G298S^ causes increased AIS length and neuronal hyperexcitability in early MNs. **A**, Schematic showing differentiation of hiPSCs into early MNs. **B**, Sanger sequencing showing CRISPR-Cas9 mediated correction of TDP-43^G298S^ mutation in hiPSCs. **C**, Reconstructions of the AIS (red) in 6-week wildtype (n=21), TDP-43^G298S^ CRISPR-corrected (n=25), TDP-43^G298S^ (n=26) MNs based on Ankyrin-G immunofluorescence staining. Counterstained against CHR2-YFP (Cyan). Scale bar = 50μm. **D**, Quantification of the average Ankyrin-G intensity profile along the AIS and total AIS length. **E**, Brightfield images showing whole-cell patch clamp recordings of 6-week CHR2-YFP positive wildtype (n=21), CRISPR corrected (n=28) and TDP-43^G298S^ (n=28) MNs. Scale bar = 50μm. **F**, Current threshold for AP firing and representative single AP traces taken from current clamp recordings. **G**, Relationship between injected current and firing frequency and maximum evoked AP firing frequency taken from current clamp recordings. **H**, Spontaneous AP firing frequency taken from passive membrane recordings of all spontaneously firing neurons. **I**, Schematic outlining main findings: TDP-43^G298S^ increases AIS length in 6-week early MN cultures. Error bars represent the SEM. p-values from one-way ANOVA tests with Dunnet’s comparison, except 1h: Mann Whitney test. *p<0.05, **p<0.01, ***p<0.001.

Spatial extension of the AIS has been shown to increase neuronal excitability by lowering the current and voltage thresholds for AP spiking (Goethals and Brette, 2020; Gulledge and Bravo, 2016; Jamann *et al*., 2021; Kole and Brette, 2018; Kuba *et al*., 2010). By performing whole-cell patch clamp recordings on early 6-week MNs we found that pathogenic TDP-43^G298S^ caused increased neuronal excitability. This was evidenced by a reduction in the current threshold for AP-spiking (Fig. 1f) as well as a shift in the input-output relationship of injected current to AP firing, and an increase in the maximum firing frequency (Fig. 1g). There was no significant change in voltage threshold (Supplementary Table 1). We also observed an increase in spontaneous firing frequency (Fig. 1h) as well as increased inward currents (most likely representing Na+) and AP amplitude (Supplementary Figure. 3b,c). These changes in excitability could not be explained by changes in passive properties (Supp. Table 1). Taken together our findings suggest that pathogenic TDP-43^G298S^ causes increased AIS length, which is linked to hyperexcitability of early 6-week hiPSC MNs (Figure 1i).

### 2. Pathogenic TDP-43^G298S^ reduces activity-dependent plasticity of the AIS in early MNs

Activity-dependent fine tuning of AIS position and/or length is a mechanism by which neurons modulate their excitability. Numerous studies have reported shortening of the AIS in response to elevated activity is associated with a compensatory reduction in excitability (Evans *et al*., 2015; Galliano et al., 2021; Jamann *et al*., 2021; Kuba *et al*., 2010; Pan-Vazquez et al., 2020; Sohn *et al*., 2019), while other studies have shown a similar homeostatic modulation of excitability associated with a distal relocation of the AIS (Grubb and Burrone, 2010; Hatch et al., 2017; Lezmy et al., 2017; Wefelmeyer *et al*., 2015). To our knowledge, AIS plasticity in human spinal MNs has not yet been investigated, nor the effect of ALS-related pathogenic TDP-43 mutations.

We found AIS length was strongly reduced in response to short-term optogenetic stimulation in early 6-week hiPSC-MNs, while AIS position was unchanged (Figure 2a,b). However, although TDP-43^G298S^ MNs displayed a shortening of the AIS, the change in length (ΔL) was less pronounced (ΔL −9.776μm ± 4.770) than in wildtype (ΔL −19.72μm ± 2.974) and corrected (ΔL −19.14μm ± 3.726) MNs. As a result, the difference in AIS length already present in baseline conditions (14.60μm ± 4.329) was even more pronounced after stimulation (23.96μm ± 4.129). In conjunction with this we found that the current threshold for AP spiking was significantly increased in the CRISPR-corrected MNs in agreement with a shortening of the AIS following optogenetic stimulation. Conversely, TDP-43^G298S^ MNs did not show a statistically significant increase in current threshold following optogenetic stimulation (Fig. 2c), even though a trend was observed. Although we did not see changes in voltage threshold, we did observe diverging voltage thresholds across genotypes following optogenetic stimulation; in this instance TDP-43^G298S^ MNs displayed a lower voltage threshold relative to CRISPR-corrected MNs, further contributing to neuronal hyperexcitability (Fig. 2c). Finally, we observed a significant downward shift in the relationship between injected current and AP spiking in the corrected MNs but not in the TDP-43^G298S^ MNs (Fig. 2d). Again, these differences in excitability could not be explained by changes in passive electrical properties (Supplementary Table 1). Taken together our findings suggest AIS plasticity is reduced in TDP-43^G298S^ MNs, a feature that further contributes to dysregulation of neuronal excitability.

**Figure 2.**
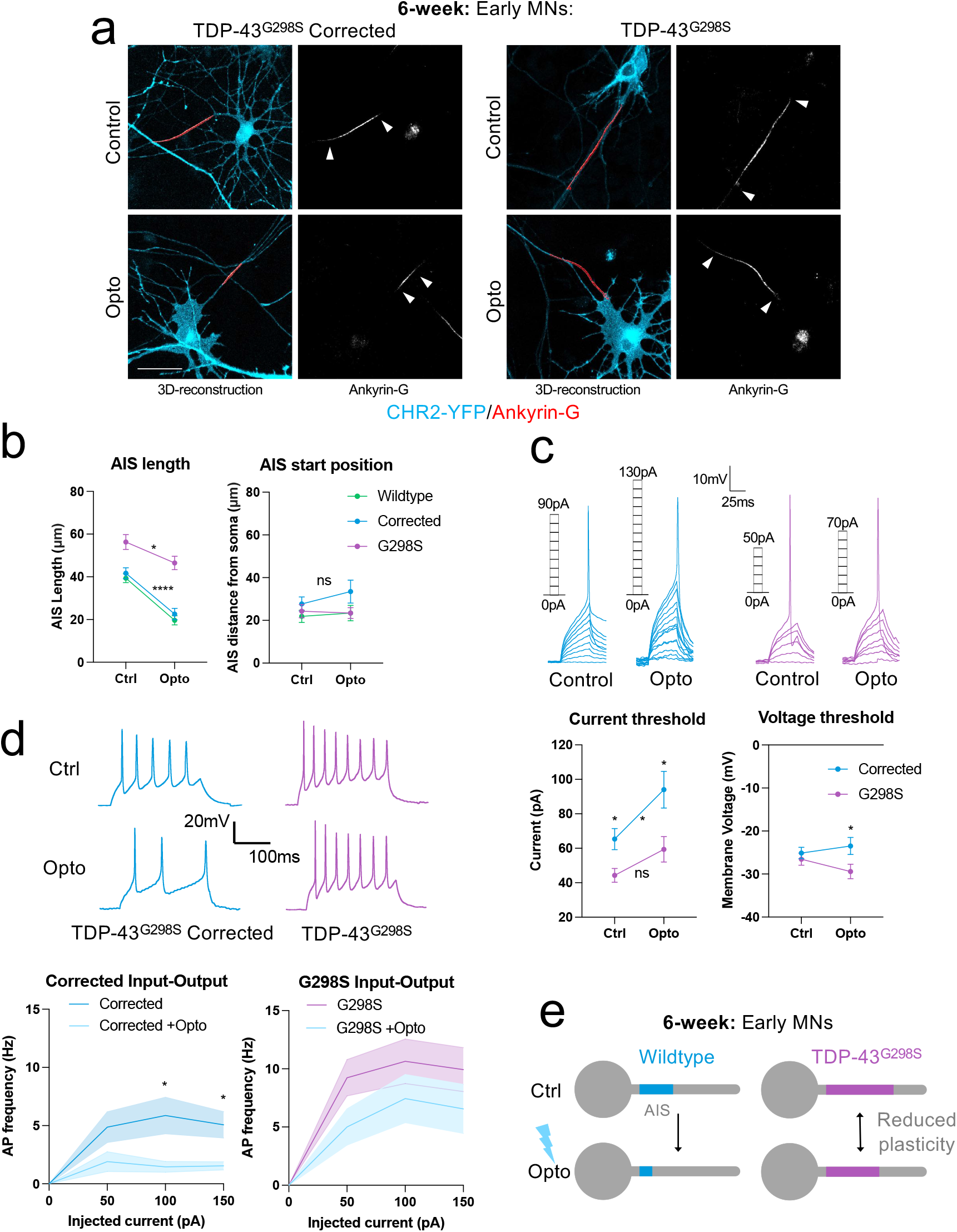
Pathogenic TDP-43^G298S^ reduces activity-dependent plasticity of the AIS in early MNs. **A**, Reconstructions of the AIS (red) in 6-week wildtype (n=21), TDP-43^G298S^ CRISPR corrected (n=25), and TDP-43^G298S^ (n=26) MNs and with 3 hours 5Hz 488nm optogenetic stimulation: wildtype (n=21), TDP-43^G298S^ CRISPR corrected (n=21), and TDP-43^G298S^ (n=21) based on Ankyrin-G immunofluorescence staining, counterstained against CHR2-YFP (cyan). Scale bar = 50μm. **B**, Quantification of AIS length change and start position in response to optogenetic stimulation. Two-way ANOVA: effect of genotype ****p<0.0001, effect of stimulation ****p<0.0001. Individual p-values from non-parametric, unpaired t-tests. **C**, Whole-cell current clamp recordings showing current and voltage thresholds for 6-week TDP-43^G298S^ CRISPR corrected (n=28), and TDP-43^G298S^ (n=28) MNs and with 3 hours 5Hz 488nm optogenetic stimulation: TDP-43^G298S^ CRISPR corrected (n=20), and TDP-43^G298S^ (n=17). Current threshold two-way ANOVA: effect of genotype **p=0.005, effect of stimulation **p=0.0046, voltage threshold two-way ANOVA: effect of genotype *p=0.025. Individual p-values from non-parametric, unpaired t-tests. Representative single AP traces and current steps. **D**, Relationship between injected current and firing frequency with and without optogenetic stimulation taken from current clamp recordings, representative AP firing traces at 100pA current injection. **E**, Schematic showing reduced activity-dependent length plasticity of the AIS in 6-week early TDP-43^G298S^ MNs. Error bars represent the SEM. *p<0.05, **p<0.01, ***p<0.001, ****p<0.0001.

### 3. hiPSC-derived neuromuscular circuits containing TDP-43^G298S^ MNs display increased spontaneous myofiber contractility

We set out to understand how changes at the AIS would impact motor unit excitability. In the early stages of ALS, motor units are hyperexcitable, leading to spontaneous muscle fasciculations (Piotrkiewicz *et al*., 2008; Shimizu *et al*., 2014). We generated an in vitro model of ALS motor units by co-culturing 6-week TDP-43^G298S^ MNs with wildtype hiPSC-myofibers in compartmentalised microdevices. Motor axons projected from outer compartments, through micro-channels to innervate myofibers in a central compartment, mimicking the spatial separation occurring in vivo (Figure. 3a). Motor unit functionality was confirmed through optogenetic stimulation of MNs, which induced robust myofiber contractions that could be abolished by the acetylcholine receptor (AChR) blocker d-tubocurarine (DTC) (Fig. 3d,f, Supplementary figure 2c).

**Figure 3.**
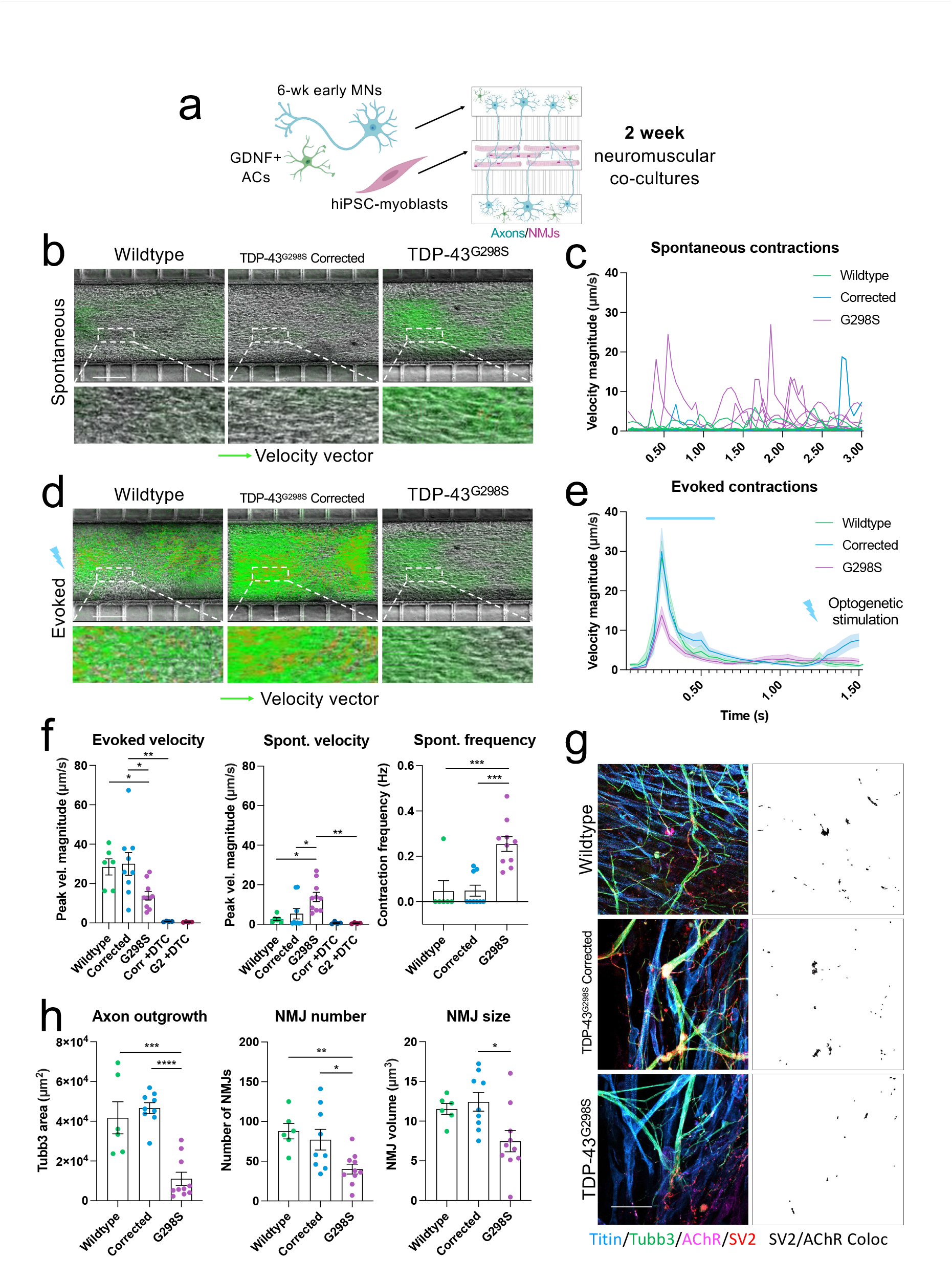
hiPSC-derived neuromuscular circuits containing TDP-43^G298S^ MNs display increased spontaneous myofiber contractility. **A**, Schematic showing generation of neuromuscular co-cultures in compartmentalised microdevices. **B**, Particle image velocimetry (PIV) analysis of spontaneous myofiber contractions co-cultures containing wildtype (n=6), CRISPR-corrected (n=9) and TDP-43^G298S^ (n=10) MNs. Velocity vectors shown in green, scale bar = 200μm. **C**, Overlayed spontaneous contraction traces in. **D**, PIV analysis of optogenetically evoked myofiber contractions. **E**, Combined traces of optogenetically evoked myofiber contractions. **F**, Quantification of spontaneous and optogenetically evoked myofiber contraction velocity with and without addition of the AChR blocker d-Tubocurarine (DTC), and quantification of spontaneous myofiber contraction frequency (Hz). **G**, Immunofluorescence staining of neuromuscular co-cultures for TTN (cyan), TUBB3 (green), SV2 (red) and AChR (magenta), and SV2/AChR colocalization channel in black. Scale bar = 50μm. **h**, Quantification based on immunofluorescence staining of axon outgrowth and neuromuscular synapse number and morphology based on SV2/AChR colocalization. Error bars represent the SEM. p-values from one-way ANOVA tests with Dunnet’s comparison: *p<0.05, **p<0.01, ***p<0.001, ****p<0.0001.

We found that co-cultures containing TDP-43^G298S^ MNs displayed increased spontaneous myofiber contraction frequency and velocity (Fig. 3b,c,f, Supplementary Movies 2,4). In contrast, optogenetically evoked contractions were weaker in the TDP-43^G298S^ co-cultures relative to the CRISPR-corrected and wildtype co-cultures (Fig. 3d,e,f, Supplementary Movies 1,3). We saw a reduction in axon coverage and neuromuscular synapses in the TDP-43^G298S^ cultures, despite axon outgrowth being initially normal (Fig. 3g,h; Supplementary Figure. 2d). This suggests that fewer neuromuscular synapses in the TDP-43^G298S^ co-cultures is responsible for the reduced maximal contractile output. Taken together this data shows that pathogenic TDP-43^G298S^ causes increased motor unit excitability, while reducing maximal contractile output by reducing the total number of functional motor units.

### 4. Pathogenic TDP-43^G298S^ causes AIS truncation, impaired plasticity, and hypoexcitability in late MNs

While neuronal hyperexcitability is a feature in the early stages of ALS, several studies have also observed a transition to hypoexcitability (Devlin *et al*., 2015; Martinez-Silva *et al*., 2018). We wanted to assess whether this occurred in our model system and whether it might be associated with changes to the AIS. We matured hiPSC-MNs for up to 10 weeks and observed a robust transition to mature repetitive firing patterns, stronger inward and outward currents and mature AP properties (Supplementary Figure. 3), as well as more mature passive membrane properties (Supplementary Table 1).

We found that pathogenic TDP-43^G298S^ caused truncation of the AIS in late MNs relative to CRISPR-corrected controls (Figure. 4a,b). This was accompanied by neuronal hypoexcitability as evidenced by a large downward shift in the input-output relationship of injected current to AP spiking and a reduction in the maximum firing frequency (Fig. 4c,d). We also observed a significant reduction in putative Na^+^ and K^+^ currents in the TDP-43^G298S^ MNs at this stage (Supplementary Figure. 3b,c). Furthermore, activity-dependent AIS length plasticity was completely abolished in TDP-43^G2398S^ MNs (Fig. 4e). While CRISPR-corrected control neurons were able to reduce the length of the AIS in response to short-term optogenetic stimulation, TDP-43^G298S^ MNs showed no reduction (Fig. 4b,e). In addition to length changes, at the later time-point MNs also showed a distal relocation of the AIS in response to optogenetic stimulation (Fig. 4f). This occurred in both TDP-43^G298S^ and CRISPR-corrected neurons. Taken together this data shows that pathogenic TDP-43^G298S^ triggers an abnormal switch from hyper to hypoexcitability, which is accompanied by truncation and impaired plasticity of the AIS. It is possible that this switch was related to a shift in the ratio of nuclear to cytoplasmic TDP-43 in the TDP-43^G298S^ MNs at this later stage (Supplementary Figure. 4b).

**Figure 4.**
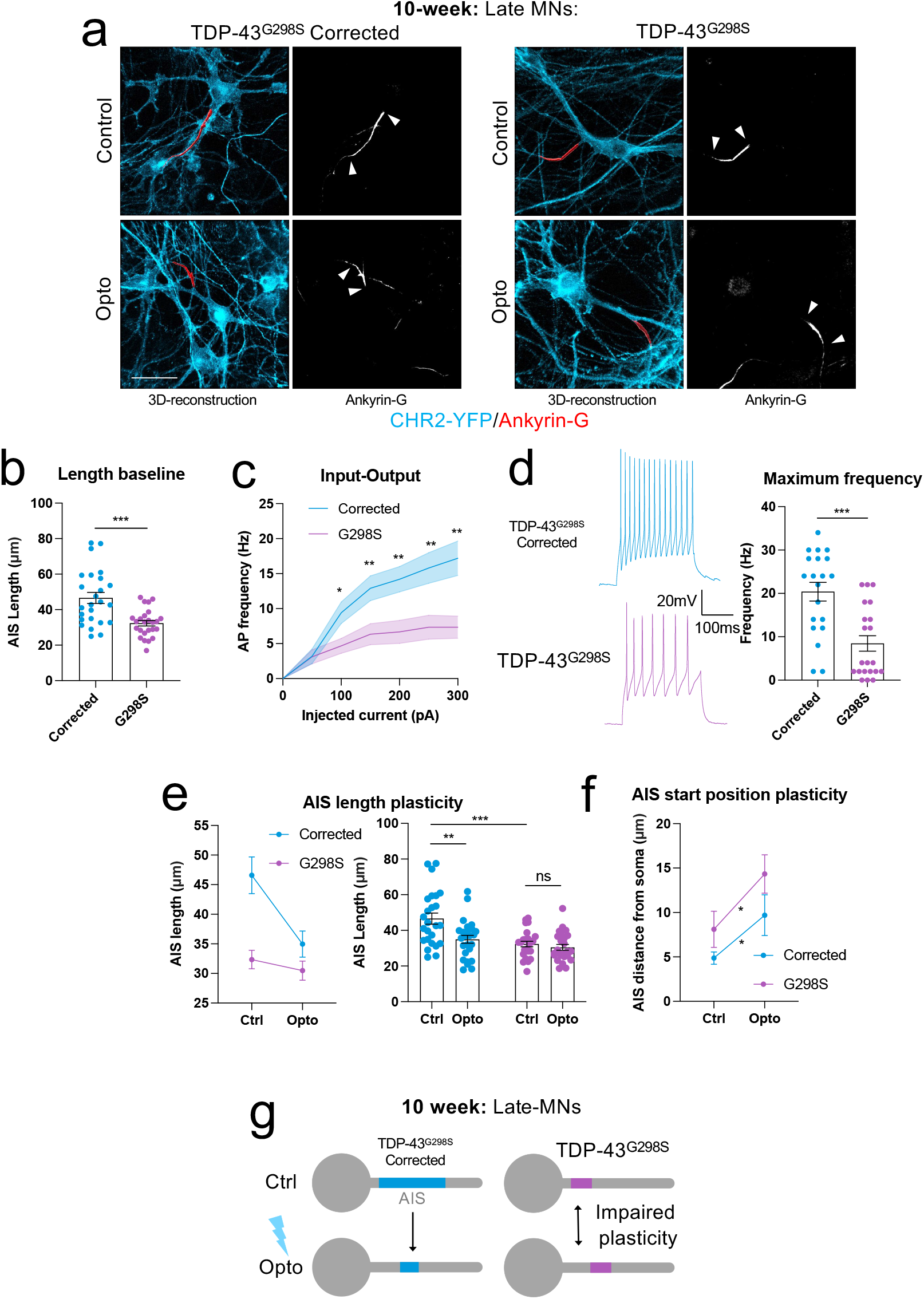
Pathogenic TDP-43^G298S^ causes AIS truncation, impaired plasticity, and hypoexcitability in late MNs. **A**, Reconstructions of the AIS (red) in 10-week wildtype (n=25), TDP-43^G298S^ CRISPR-corrected (n=25), and TDP-43^G298S^ (n=25) MNs with and without 3 hours 5Hz optogenetic stimulation. Based on Ankyrin-G immunofluorescence staining, counterstained against CHR2-YFP (cyan). Scale bar = 50μm. **B**, Quantification of AIS length based on Ankyrin-G staining in baseline unstimulated conditions. **C**, Relationship between injected current and firing frequency taken from whole-cell current clamp recordings of 10-week TDP-43^G298S^ CRISPR-corrected (n=20), and TDP-43^G298S^ (n=21) MNs. **D**, Maximum evoked AP firing frequency (Hz) taken from current clamp recordings. **E**, Quantification of AIS length change in response to optogenetic stimulation. Two-way ANOVA: effect of genotype *p<0.04, effect of stimulation **p<0.004, interaction *p<0.029. **F**, Quantification of AIS start position change in response to optogenetic stimulation, measured as start position from the soma. **G**, Schematic showing truncation and impaired plasticity of the AIS in 10-week late TDP-43^G298S^ MNs. Error bars represent the SEM. p-values from non-parametric, unpaired t-tests:*p<0.05, **p<0.01, ***p<0.001.

### 5. Pathogenic TDP-43^G298S^ drives aberrant expression of AIS genes in early and late MNs

Since we had observed dynamic changes to the structure of the AIS in early and late MNs, we sought to investigate whether altered expression of AIS specific genes could account for these changes. Indeed, TDP-43 is known to play a pivotal role in global gene expression, RNA splicing, as well as RNA stability and transport (Alami et al., 2014; Fukushima et al., 2019; Lagier-Tourenne et al., 2012; Tank et al., 2018) and a number of studies have found evidence that TDP-43 can bind and modulate splicing of a number of key AIS genes including Ankyrin-G (Brown et al., 2022; Fukushima *et al*., 2019; Lagier-Tourenne *et al*., 2012; Ma et al., 2022; Narayanan *et al*., 2013). We found that pathogenic TDP-43^G298S^ caused an increase in the expression of several AIS related genes in early (6-week) MNs (Figure 5a and Supplementary Figure 5a), including the AIS master scaffolding protein ANK3 (Ankyrin-G) as well the voltage gated sodium channels SCN1A (Nav1.1) and SCN8A (Nav1.6) responsible for AP initiation (Figure. 5a). Moreover, we found an increase in the ratio of the AIS-specific 480kDa Ankyrin-G isoform relative to the shorter non-AIS 270kDa isoform (Figure. 5b). This change in expression pattern was also associated with increased Ankyrin-G protein at the AIS (Supplementary figure. 4c). At the later (10-week) timepoint pathogenic TDP-43^G298S^ caused a decrease in overall ANK3, SCN1A and SCN8A expression (Figure. 5c) as well as an increase in the ratio of the short non-AIS 190kDa isoform relative to the 480kDa AIS specific isoform of Ankyrin-G (Figure. 5d). This change was associated with reduced Ankyrin-G protein at the AIS (Supplementary Figure. 5c). Taken together these results show that pathogenic TDP-43^G298S^ causes dysregulated expression of AIS specific genes, correlating with the structural and electrophysiological changes observed at the AIS in early and late MNs.

**Figure 5.**
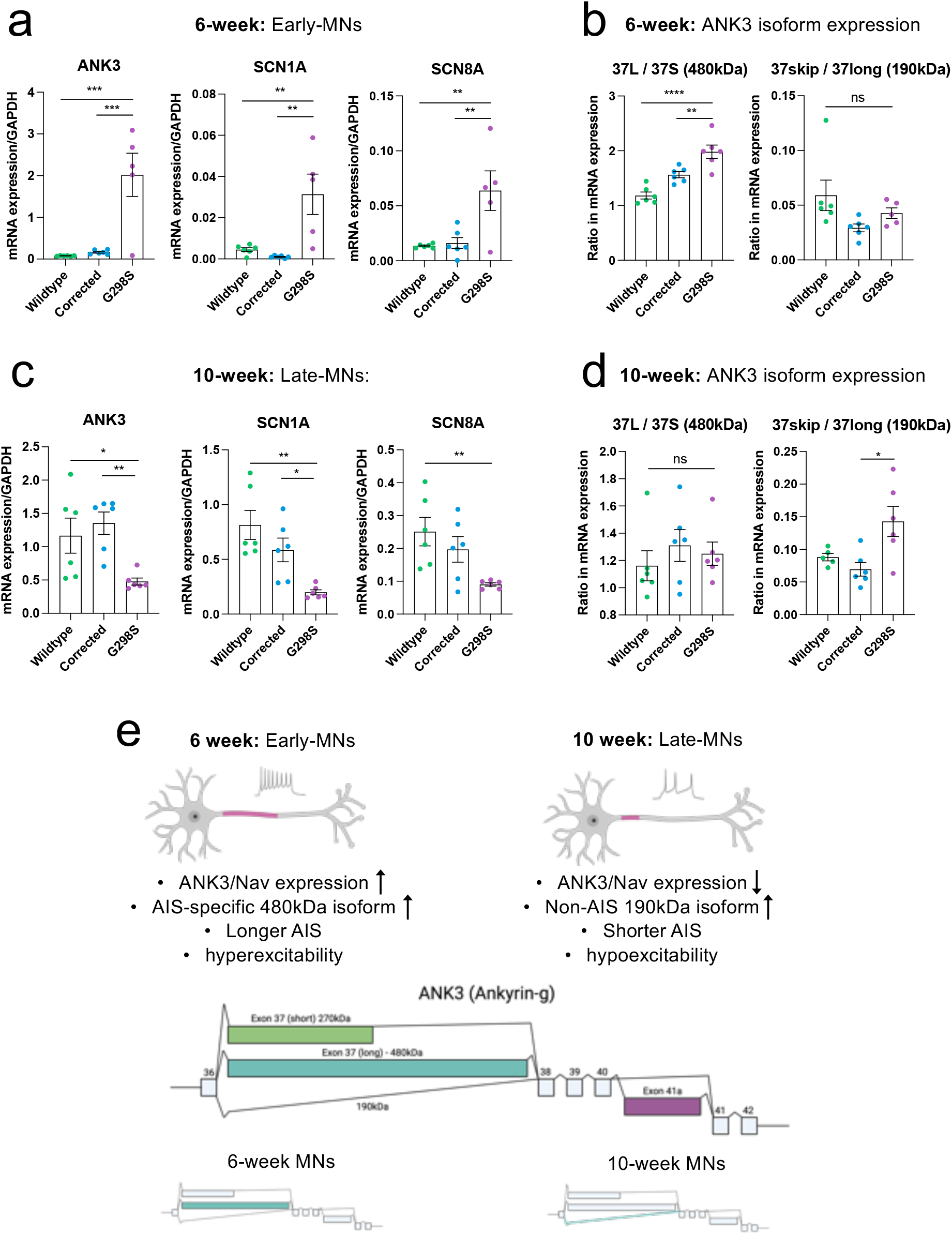
Pathogenic TDP-43^G298S^ drives aberrant expression of AIS genes in early and late MNs. **A**, qRT-PCR analysis of ANK3 (Ankyrin-G), SCN1A (Nav1.1) and SCN8A (Nav1.6) expression in early (6-week) MNs. **B**, qRT-PCR analysis of ANK3 480kDa/190kDa isoform ratios in early MNs. 37 long (480kDa) relative to 37S (270kDa), and 37 skipping (190kDa) relative to 37 long (480kDa). **C**, qRT-PCR analysis of ANK3, SCN1A, and SCN8A expression in late (10-week) MNs. **D**, qRT-PCR analysis of ANK3 480kDa/190kDa isoform ratios in late MNs. isoform expression, SCN1A, SCN2A, SCN8A, and KCNA1 expression in late (10-week) MNs. **E**, Schematic showing changes to AIS gene expression, ANK3 isoform expression, AIS structure and neuronal excitability in early and late MNs. Error bars represent the SEM. p-values from one-way ANOVA tests with Dunnet’s comparison. *p<0.05, **p<0.01, ***p<0.001, ****p<0.0001.

## Discussion

Here we report a new pathophysiological mechanism driving dysregulated neuronal excitability in human ALS MNs. We found that the ALS-related pathogenic TDP-43^G298S^ mutation caused abnormal expression of AIS specific genes, including the scaffolding protein Ankyrin-G (ANK3) and the voltage gated sodium channels Nav1.1 (SCN1A) and Nav1.6 (SCN8A), leading to structural changes at the AIS - the site of action potential generation in neurons as well as changes to neuronal excitability. Early TDP-43^G298S^ MNs showed increased expression of Ankyrin-G, leading to increased Ankyrin-G protein at the AIS, increased AIS length and neuronal hyperexcitability. Furthermore, when co-cultured with hiPSC-myofibers in compartmentalized microdevices, TDP-43^G298S^ MNs triggered spontaneous myofiber contractions, mirroring muscle fasciculations observed in patients with ALS. In contrast, late TDP-43^G298S^ MNs showed reduced expression of Ankyrin-G, leading to reduced Ankyrin-G protein at the AIS, reduced AIS length and neuronal hypoexcitability. Finally at all stages, pathogenic TDP-43^G298S^ compromised activity dependent plasticity of the AIS, further contributing to dysregulated neuronal excitability. Taken together these results point toward the AIS as an important subcellular target driving excitability changes in ALS MNs.

Structural plasticity of the AIS plays an integral role in maintaining normal levels of neuronal excitability. Previous studies have implicated the AIS as a possible target in ALS. Significant swelling of the AIS has been reported in electron microscopy studies of post-mortem ALS lumbar spinal MNs (Sasaki and Maruyama, 1992) and studies in SOD1 mice have shown altered AIS length in pre-symptomatic and symptomatic stages of the disease (Bonnevie et al., 2020; Jorgensen et al., 2021). Interestingly Jorgensen and colleagues (Jorgensen *et al*., 2021) found that extension of the AIS at symptom onset in SOD1 mice was associated with changes in Na+ currents. However, since patients with SOD1 mutations do not exhibit TDP-43 pathology (Mackenzie et al., 2007), a feature seen in the majority of ALS patients (Neumann *et al*., 2006), we wanted to investigate how pathogenic TDP-43 mutations would impact both AIS structure and homeostatic plasticity in a representative human model of the disease.

We found that pathogenic TDP-43^G298S^ caused increased length of the AIS in early hiPSC-MNs (Figure. 1c,d). This increased length was associated with stronger Na+ currents and AP amplitude (Supplementary Figure. 2d), and neuronal hyperexcitability, characterized by a shift in the input-output relationship of injected current to AP spiking, and a reduction in the current threshold for AP firing (Figure. 1f,g). Taken together with findings by Jorgensen (Jorgensen *et al*., 2021) and Sasaki (Sasaki and Maruyama, 1992) our findings implicate increased AIS size as a novel pathophysiological mechanism for modulating neuronal excitability across divergent subsets of ALS. In addition to structural changes at the AIS, we also observed reduced activity dependent plasticity of the AIS, particularly in late MN cultures (Figure. 2b, Figure. 4e). This finding has not previously been reported and has significant implications since activity-dependent fine-tuning of AIS length is an important homeostatic mechanism to prevent abnormal levels of network activity in neurons (Evans *et al*., 2015; Grubb and Burrone, 2010; Jamann *et al*., 2021; Kuba *et al*., 2010; Sohn *et al*., 2019; Wefelmeyer *et al*., 2015).

Several studies have reported hyperexcitability of hiPSC derived MNs harbouring ALS-related mutations (Devlin *et al*., 2015; Wainger et al., 2014) as well as mouse models (Jorgensen *et al*., 2021; Pambo-Pambo et al., 2009) and patient studies (Shimizu *et al*., 2014). Our study is the first to link pathogenic TDP-43 to structural and functional changes at the AIS and downstream changes in neuronal excitability. By using CRISPR-Cas9 gene correction to rescue these phenotypes we can be confident that the TDP-43^G298S^ mutation accounts for these changes rather than variability between hiPSC-lines (Figure. 1b; Supplementary Figure 1a). Furthermore, we generated synchronized cultures of highly MACS enriched MNs to prevent inadvertent recordings from immature progenitors and unrelated contaminating neuronal subtypes (Supplementary Figure. 1d). By using wildtype astrocytes and myofibers in our co-cultures, we can be confident that the effects we observe are cell autonomous.

Neuronal hyperexcitability has long been suspected of contributing to excitotoxicity and MN vulnerability in ALS. Indeed, one of the few approved drugs to extend survival in ALS, Riluzole, primarily acts to dampen neuronal excitability by inhibiting persistent inward Na+ currents as well as dampening glutamate induced excitotoxicity (Doble, 1996; Song et al., 1997). Furthermore, the incidence of peripheral axonal and motor unit hyperexcitability leading to complex muscle fasiculations has been shown to correlate with increased disease severity and reduced survival time in patients with ALS, suggesting that this process influences progression of the disease. This led us to investigate how TDP-43-dependent alterations to AIS structure and function would impact motor unit excitability. Based on our previously published work (Machado *et al*., 2019; Paredes-Redondo et al., 2021) and other similar models (Osaki et al., 2018; Rao et al., 2018) we generated a model of ALS motor units by co-culturing hiPSC-MNs and wildtype iPAX7 hiPSC-myofibers in compartmentalized microfluidic devices (Figure. 3a). We found that neuromuscular circuits containing TDP-43^G298S^ MNs showed an increase in spontaneous myofiber contractions (Figure 3.b,c,f,g). Conversely these cultures showed a reduction in the maximal evoked contractile output, (Figure. 3d,e,f) a finding linked to a reduction in the total number of neuromuscular synapses (Figure. 3h,i). The combination of these two phenotypes closely resembles the simultaneous occurrence of muscle fasciculations and muscle weakness that occurs in patients with ALS (Hardiman *et al*., 2017; Shimizu *et al*., 2014). We believe this platform has great potential for developing drugs to target these common features of ALS.

In later MNs we observed a switch from early hyperexcitability to hypoexcitability (Figure. 4; Supplementary Figure 2d). Hypoexcitability has previously been reported in hiPSC models of ALS (Naujock et al., 2016; Sareen et al., 2013), and a switch from hyper to hypoexcitability has also been demonstrated in hiPSC-MNs harbouring TDP-43 mutations and C9orf72 hexonucleotide expansions (Devlin *et al*., 2015) and in the SOD1 mouse model (Martinez-Silva *et al*., 2018). Here, we found that this switch was linked to structural changes and impaired activity-dependent plasticity of the AIS (Figure. 4c,f,g). Specifically, we observed truncation of the AIS and complete abolishment of activity dependent AIS length plasticity at this stage. We also observed a large reduction in putative Na+ and K+ currents (Supplementary Figure. 3b,c).

Finally, we uncovered a molecular basis explaining the switch from early AIS extension and hyperexcitability to later AIS truncation and hypoexcitability. We found that early TDP-43^G298S^ MNs showed increased expression of the AIS master scaffolding protein Ankyrin-G (ANK3), as well increased expression of the voltage gated ion channels Nav1.1 (SCN1A) and Nav1.6 (SCN8A) that are important for AP initiation. Furthermore, we also observed an increase in the ratio of the long 480kDa AIS specific isoform of Ankyrin-G relative to shorter non-AIS isoforms. In conjunction with this we observed a corresponding increase Ankyrin-G protein at the AIS which could directly account for the increase in AIS length and hyperexcitability observed at this stage. In contrast we found that late TDP-43 MNs expressed less Ankyrin-G, Nav1.1 and Nav1.6 and showed an increase in the ratio of the short non-AIS 190kDa Ankyrin-G isoform that typically localizes to synapses relative to the longer 480kDa AIS isoform. This was associated with reduced Ankyrin-G at the AIS, reduced AIS length and neuronal hypoexcitability. Indeed, several studies have identified Ankyrin-G as a downstream target of TDP-43, indicating that our observed data could be a direct result of abnormal TDP-43 regulation of Ankyrin-G expression and splicing (Lagier-Tourenne *et al*., 2012; Ma *et al*., 2022; Narayanan *et al*., 2013). It’s also possible that TDP-43 modifies expression of upstream transcription factors that control larger networks of AIS-specific genes such as RBFOX (Jacko et al., 2018). Alternatively, it is possible that TDP-43 differentially regulates AIS mRNA transcript stability, degradation and transport in the early and late MN cultures (Fukushima *et al*., 2019). Interestingly it was only at the later timepoint we observed direct evidence of TDP-43 mis-localisation, possibly accounting in the switch in phenotypes observed. The early phenotype may be related specifically to the mutant TDP-43 protein, while the later switch in phenotype may be a result of TDP-43 mis-localisation that takes longer to develop, and perhaps more representative of a later disease stage.

In summary, we report that the ALS-related pathogenic TDP-43^G298S^ mutation causes abnormal expression of AIS specific genes including the scaffolding protein Ankyrin-G, leading to disrupted structure and activity-dependent plasticity of the AIS in hiPSC-MNs. In early neurons increased expression Ankyrin-G led to increased AIS length, neuronal hyperexcitability, and increased spontaneous myofiber contractions in compartmentalised neuromuscular co-cultures. Conversely in late MNs reduced expression of Ankyryin-G was associated with reduced AIS length and hypoexcitability. Taken together these results highlight the AIS as an important target driving excitability changes in ALS.

---

**Supplementary Movie 1**. TDP-43^G298S^ CRISPR corrected evoked contraction

**Supplementary Movie 2**. TDP-43^G298S^ CRISPR corrected spontaneous contraction

**Supplementary Movie 3**. TDP-43^G298S^ Evoked contraction

**Supplementary Movie 4**. TDP-43^G298S^ Spontaneous contractions

## Acknowledgments

We would like to thank Siddharthan Chandran, Agnes Nishimura and Christopher Shaw for sharing cell lines and reagents. Oliver Baker, Ieva Berzanskyte, Benjamin Compans, Vincenzo Mastrolia, Winnie Wefelmeyer for advice on experimental techniques. Winnie Wefelmeyer, Matthew Grubb, Benedikt Berninger, Nicolas Marichal Negrin, and Geraldine Jowett for comments on the manuscript. The authors acknowledge financial support from the Medical Research Council (Grant No. MR/N025865/1) to IL, the Wellcome trust (Investigator award 215508/Z/19/Z) and BBSRC (Grant No. BB/S000526/1) to JB. PH and FR were supported by Wellcome Trust “Cell Therapies & Regenerative Medicine” PhD studentships (108874/Z/15/Z), and AC by a BBSRC LIDo PhD studentship (BB/M009513/1). The authors also acknowledge financial support from the Department of Health via the National Institute for Health Research (NIHR) comprehensive Biomedical Research Centre award to Guy’s & St Thomas’ NHS Foundation Trust in partnership with King’s College London and King’s College Hospital NHS Foundation Trust. The authors acknowledge the Medical Research Council Centre grant MR/N026063/1.

This research was funded, in part, by the Wellcome Trust [215508/Z/19/Z]. For the purpose of open access, the author has applied a CC BY public copyright licence to any Author Accepted Manuscript version arising from this submission.

## Author contributions

Conceptualisation by PH, JB, IL. Study design and methodology by PH, IL, JB, GN, FR, CBM. Experimental work was carried out by PH, FR, CBM, AC, LRB, IL. Data analysis carried out by PH, GN. Original draft by PH, GN, JB, IL. Review and editing by PH, GN, JB, IL.

## Methods

### Experimental model and subject details

#### Engineering hiPSC-lines

Patient-derived hiPSC line harbouring the pathogenic TDP-43^G298S^ mutation was provided by Christopher Shaw (King’s College London) and Siddarthan Chandran (The University of Edinburgh). The Line was originally published in: (Barton et al., 2021). Wildtype hESC H9 line was acquired from WiCell (Madison. Wisconsin, USA) under a license from the steering committee for the UK Stem Cell Bank (No. SCS11-06). HipSci Lines samples were collected from consented research volunteer recruited from the NIHR Cambridge BioResource through (http://www.cambridgebioresource.org.uk). Initially, 250 normal samples were collected under ethics for iPSC derivation (REC Ref: 09/H0304/77, V2 04/01/2013), which require managed data access for all genetically identifying data, including genotypes, sequence and microarray data (‘managed access samples’). In parallel the HipSci consortium obtained new ethics approval for a revised consent (REC Ref: 09/H0304/77, V3 15/03/2013), under which all data, except from the Y chromosome from males, can be made openly available (Y chromosome data can be used to de-identify men by surname matching), and samples since October 2013 have been collected with this revised consent (‘open access samples’).

hiPSC cell clones intended for MN differentiation and enrichment were engineered to express the HB9::hCD14 MACS sortable construct using TALENS based insertion into the AAVS1 safe-harbour locus (SFigure. 1d) (Hockemeyer et al., 2009) using a NEPA-21 electroporator (Sonidel). Cell lines were also engineered to express the optogenetic actuator transgene CAG::CHR2-YFP using a PiggyBAC-mediated integration system via electroporation. Fluorescence activated cell-sorting (FACS) was carried out using a BD FACSAria™ 3 (BDBiosciences) to select CHR2-YFP positive cells to generate polyclonal cell lines with comparable YFP expression (SFigure. 1e). Inducible iPAX7 hiPSC-lines to forward program hiPSCs into myoblast progenitors were generated by TALENS based integration the Doxycycline-inducible PAX7 construct into the CLYBL safe-harbour locus (Cerbini et al., 2015) of the publicly available HiPSCi line PAMV1 (www.hiPSCi.org), via electroporation.

#### Cell culture and differentiation

hiPSCs/ESCs were maintained on 0.4μg/cm^2^ LN521 basement matrix (BioLamina) in StemMACS iPS-brew XF with iPSbrew supplement (1X) (Miltenyi Biotec) plus Penicillin/streptomycin (1x) (Gibco). Cells were passaged at 70% confluency as single cells by incubating cells with TrypLE express (Invitrogen) for 5 min at 37°C and replating in 10μM ROCK inhibitor Y-27632 (Tocris) for 24h.

MN differentiation was based on (Du et al., 2015) with minor modifications. hiPSCs were grown until 100% confluent on 0.4μg/cm^2^ LN521 basement matrix (BioLamina) in StemMACS iPS-brew XF with iPSbrew supplement (1X) (Miltenyi Biotec). Cells were then passaged 1:3 as clumps using 15 min 37°C incubation with 1mg/ml collagenase IV (Invitrogen) onto plates coated with 1:50 GFR-Matrigel (Corning) in iPS-brew XF containing 10μM ROCK inhibitor. After 2-3 days once colonies started to merge media was switched to NEP media (d0), comprising basal media: 1-part DMEM/F-12 (Gibco), 1-part Neurobasal (Gibco), N2 supplement (0.5x) (Gibco), Neurobrew-21 (0.5x) (Miltenyi Biotec), L-Glutamine (2mM)

(Gibco), Penicillin/streptomycin (1x) (Gibco), plus 3μM CHIR99021 (Tocris), 2μM SB431542 (Tocris) and 0.2μM LDN193189 (Stemgent) for 6 days. After 6 days cells were again split 1:3 with collagenase IV onto 1:50 matrigel coated plates and media changed to MNP media, comprising basal media plus 0.1μM RA (Sigma), 0.5μM purmorphamine (Tocris), 1μM CHIR99021, 2μM SB431542, 0.2μM LDN193189. After a further 6 days cells were either frozen or split 1:3 using collagenase IV and switched to MNP expansion media, comprising basal media plus 0.1μM RA, 0.5μM purmorphamine, 3μM CHIR99021, 2μM SB431542, 0.2μM LDN193189, 0.5mM valproic acid (Stemgent). Cells were passaged up to 2 times in expansion conditions before freezing.

Myoblast differentiation was based on (Rao *et al*., 2018) with minor modifications, using the custom built iPAX7 knock-in hiPSC line rather than lentiviral transduction. At 70% confluency hiPSCs were split onto 1:50 Matrigel coated plates at a density of 33k/cm^2^ in iPS-brew XF plus 10μM ROCK inhibitor Y-27632. The following day media was replaced with E6 media (Gibco) plus L-Glutamine (2mM) and Penicillin/streptomycin and 10μM CHIR99021 for 2 days, after which CHIR99021 was removed and replaced with 1 µg/mL Doxycycline (Sigma) for 18 days with 10ng/ml bFGF (R&D) added from day 5. Myoblast progenitors were cultured on 1:50 matrigel coated flasks in expansion media, comprising low glucose DMEM (ThermoFisher), with 10% FBS (Gibco), 1x NEAA (Gibco), L-Glutamine (2mM), Penicillin/streptomycin, 1x β-Mercaptoethanol (Gibco) and expanded up to 4 passages. To differentiate the myoblasts into myotubes cells were seeded on 1:50 Matrigel coated plates at a density of 100k/cm^2^ into myogenic differentiation media, comprising low glucose DMEM, 0.5x N2 supplement, 10% horse serum (Gibco), L-Glutamine (2mM), 1x Penicillin/streptomycin.

GDNF+ mESC astrocyte differentiation was performed as described in (Machado *et al*., 2019).

### Method details

#### Magnetic-activated cell sorting (MACS)

Motor neuron progenitors (MNPs) were thawed in MNP differentiation medium, comprising basal medium plus 0.5μM RA, 0.1μM purmorphamine and ROCK inhibitor Y-27632 for 24h at density of 600k/cm^2^ onto Matrigel coated plates. Cells were grown in these conditions for 1 week prior to MACS sorting. Cells were then dissociated in TrypLE +10U/ml DNase (Roche) for 5-7 mins at 37°C into a single cell solution then washed 3x in DMEM +10U/ml DNase. Cells were then filtered using a 40µm nylon strainer (BD Falcon) and re-suspended in MACS buffer comprising PBS (Gibco), 10% BSA (ThermoFisher), +10U/ml DNase and 3μg/ml anti-CD14 antibody (clone 26ic, DSHB), and transferred to a MACSmix tube rotator (Miltenyi Biotec) at 4°C for 15 min. Cells were then washed and resuspended in MACS buffer plus 1:5 anti-mouse igG microbeads (Miltenyi Biotec) and rotated at 4°C for a further 15 min. Cells were then resuspended in 1ml MACS buffer and applied to a MS magnetic column mounted to an OctoMACS magnet (Miltenyi Biotec). Cells were washed 3x with 500μl MACS buffer then the positive fraction eluted in 1ml MACS buffer.

#### Targeted gene editing

Targeted gene editing in hiPSCs was achieved using CRISPR-Cas9 mediated homology directed repair to correct the endogenous TDP-43^G298S^ mutation. 3μg TrueCut CAS9 v2 (ThermoFisher) was combined with 90μM sygRNA (5’ - GGATTTGGTAATAGCAG AGGGGG 3’) (Merck) at RT for 15 minutes to form an RNP complex. This complex was then electroporated using a NEPA-21 electroporator (Nepagene) into 1×10^6^ hiPSCs along with 50μM ddOligo template DNA (ThermoFisher). The donor template was engineered with the GGT codon to correct the AGT codon responsible for the TDP-43^G298S^ mutation and also with a silent Xho1 restriction site in the seed region of the sgRNA. Clones were then screened for the presence of the Xho1 restriction site and positive clones sequenced using sanger sequencing (SourceBioscience). Subsequently we sequenced ∼1000bp upstream and downstream of exon 6 of TARDBP and the top 5 predicted off-target sgRNA binding sites to confirm no off-target genome editing. We then carried out g-banding to confirm a normal karyotype (CellGuidanceSystems) (SFigure. 1b), and Western blot analysis of TDP-43 using an n-terminal TDP-43 antibody (MAB7778 - Biotechne) to confirm normal protein expression (SFigure. 1c).

#### Immunofluorescence

For imaging, 5k MACS sorted MNs were seeded onto 20k MACS sorted GDNF+ mESC-derived astrocytes in 96-well high content imaging plates (Greiner 655090) and grown in MN maturation medium, comprising basal medium plus 0.1μM RA, 0.1μM purmorphamine and ROCK inhibitor Y-27632. Cells were then fixed in 4% PFA for 15 min at RT and washed 3x in PBS. Subsequently cells were blocked in 3% BSA and 0.1% Triton X-100 in PBS for 1hr at RT. Neuromuscular co-cultures were additionally permeabilised using 10% DMSO. Cells were incubated overnight with mouse IgG2a anti-AnkG (N106/36 NeuroMab), rabbit anti-GFP (A11122 Invitrogen), rabbit anti-isl1 (ab109517 Abcam), mouse IgG2a anti-Tubb3 Tuj1 (MAB1995 R&D systems), mouse IgG1 anti-SV2 (DSHB), rat anti-Mab35 (DSHB) and mouse igM anti-Titin (9D10 DSHB) in blocking buffer at 4°C. Cells were then washed 3x in 0.1% Triton X-100 in PBS and incubated with the secondary antibodies: anti-mouse IgG2a 647 (Alexa Fluor R&D systems), anti-mouse IgG2a 488 (Alexa Fluor R&D systems), anti-rabbit 488 (Alexa Fluor R&D systems) anti-rabbit 647 (Alexa Fluor R&D systems), anti-mouse IgM 405 (Alexa Fluor R&D systems), anti-mouse IgG1 647 (Alexa Fluor R&D systems, anti-rat 555 (Alexa Fluor R&D systems) and DAPI (Sigma) for 2hr at RT. Cells were final washed 3x in PBS. Cells were then imaged using a Leica TCS SP8 confocal inverted laser scanning microscope at a 63x oil objective.

#### qRT-PCR

Total RNA was extracted using a pure link RNA kit (Invitrogen) according to manufacturer’s instructions, and subsequently reverse transcription was performed using a GoScript reaction kit (Promega) to produce cDNA. 5ng of cDNA was loaded per qRT-PCR reaction using fast SYBR master mix and reactions run on a CFX96 RT-PCR detection system (Bio-Rad). The following primers were used:

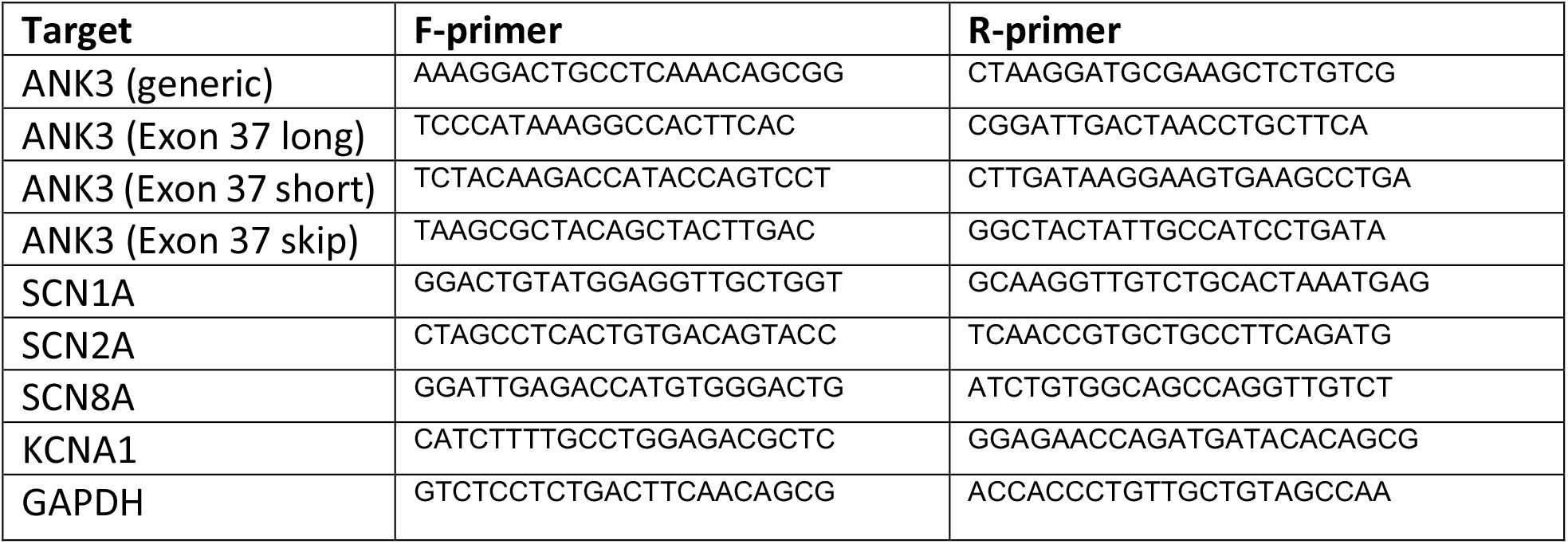

#### Electrophysiology

For patch clamp recordings 50k MACS sorted MNs and 50k MACS sorted mESC-GDNF astrocytes were seeded onto 1:50 Matrigel coated 18mm glass coverslips and cultured in MN maturation medium, comprising basal medium plus 0.1μM RA, 0.1μM purmorphamine and ROCK inhibitor Y-27632. Coverslips were transferred to an open bath chamber (RC-41LP Warner Instruments) containing extracellular solution: NaCl 136 mM, KCl 2.5 mM, HEPES 10 mM, MgCl2 1.3 mM, CaCl2 2 mM, Glucose 10 mM, pH adjusted to 7.3 and osmolarity adjusted to 300 mOsm. The chamber was mounted on an inverted epifluorescence microscope (Olympus IX71) and visualised using a 60x oil objective. Pipettes were pulled from borosilicate glass (O.D. 1.5mm, I.D. 0.86mm, Sutter instruments) to a resistance of between 3-5MΩ. Intracellular solution contained the following: 125 mM KMeSO4, 5 mM MgCl2, 10 mM EGTA, 10 mM HEPES, 0.5 mM NaGTP, 5 mM Na2ATP, pH = 7.4, osmolarity = 290mOsm. Whole-cell patch clamp recordings were then made at the soma of CHR2-YFP+ MNs using a Multiclamp 700B amplifier (Molecular Devices) and the data acquired using a Digidata 1440A digitizer (Molecular Devices). All recordings were carried out at room temperature. Data was acquired with Clampex software (Molecular Devices) and Axon Multiclamp Commander Software (Molecular Devices). Current-clamp data was sampled at a rate of 50kHz and filtered at 10kHz and Volatge-clamp data was sampled at 20 kHz and filtered at 10 kHz. Whole-cell currents used to estimate Na^+^ and K^+^ conductances were recorded in voltage clamp using 50 ms voltage steps from −80mV to +50mV. Resting membrane potentials and spontaneous AP spiking were recorded for 1 minute in current clamp mode without current injection, within the first 2 minutes after break-in. Intrinsic excitability measurements and AP properties were recorded in current clamp mode, while using a steady current injection to maintain membrane potential close to −60 mV (values for injected current and membrane voltage were indistinguishable across genotypes and are reported on Supplementary Figure 4).using either 100 ms current injections from −20pA – 170pA (for measurements of AP properties) or 500ms current injections from −50pA to 300pA (for measurements of Input-output characteristics). Although typically shorter current steps are used for establishing AP properties (eg - current threshold), these were unreliable at eliciting APs in younger MNs (6 week cultures). We therefore used 100ms current steps instead, which allowed us to compare AP properties across all stages. Assessment of optogenetic stimulation was carried out by delivering 500 ms pulses of 488 nm light were applied using a CoolLED pE-100 illumination system and AP traces recorded in current clamp with membrane potential set to −60 mV.

#### Microdevice fabrication and neuromuscular co-cultures

Microdevices were made according to our previously published work (Machado *et al*., 2019) with slight modifications. A thin layer of NOA-73 resin (Norland products) was applied to plastic bottom dishes (diameter, 35 mm; ibidi, 81156) with a cell scraper and partially UV– cured for 10 s at 55 J/cm2. Then, the PDMS arrays were placed on the resin and fully UV-cured for 1 min. Following UV-sterilization of the devices, two neural spheroids comprising 10k MACS sorted MNs and 5k MACS sorted astrocytes formed in lipidure coated (Amsbio) U-bottom 96 well plates were loaded into each of the two outer-compartments. Co-cultures were optogenetically entrained using 5Hz 450nm optogenetic stimulation for 1hr a day for 4 days, 3 days after plating 20k myoblasts in the central compartment, in order to enhance neuromuscular synapse formation.

#### Optogenetic stimulation

To analyse changes to the AIS and electrophysiological parameters in response to short term activity we optogenetically stimulated cultures using a custom-built heat sink and LED assembly (Machado *et al*., 2019). Custom written software controlled the timing of LED emission. LEDs emitted pulses of 450nm blue light at a frequency of 5Hz with a 20ms epoch set at an LED intensity of 40%. Cultures were supplemented with 1x antioxidant supplement (Sigma) to mitigate the effects of phototoxicity.

### Quantification and Statistical analysis

#### Analysis of AIS morphology

AIS length, diameter and start position were analysed using FIJI. Ankyrin-G fluorescence was uniformly thresholded and the AIS length and position relative to the soma, as determined by the CHR2-YFP counter stain, were traced. AIS diameter was measured at the mid-point of the AIS. Reconstructions of the AIS were carried out using IMARIS 9.1.2 software by generating a surface map based on the Ankyrin-G fluorescence. Gating thresholds for this were kept constant between samples.

#### Analysis of axon outgrowth and detection of NMJs

Axon outgrowth and neuromuscular junctions were analysed using IMARIS 9.1.2 software. For axon outgrowth, 3D surface maps were generated based on TUBB3 immunofluorescence using uniform gating thresholds. From these reconstructions the total axon volume and surface area per field of view were calculated. For analysis of neuromuscular junctions, the IMARIS coloc function was used to automatically generate a colocalization channel for the pre-synaptic SV2 immunofluorescence channel and the post-synaptic AChR channel. 3D surface maps were then generated based on this coloc channel using uniform gating thresholds. From this surface map the total number of colocalised objects as well as the volume and surface area of these objects could be calculated.

#### Particle image velocimetry (PIV) analysis of myofiber contractions

Video recordings of spontaneous and optogenetically evoked myofiber contractions were analysed by Particle Image Velocimetry using the PIVlab package in Matlab. Three iterations of interrogation windows of 64/32/16 pixels, each with 50% overlap were used and frames calibrated to a known reference distance. Vectors were then validated by filtering out velocity values higher than 7 times the standard deviation. Mean velocity values for the myofiber compartment area were exported to derive peak velocity magnitude values and contraction frequency values. To show that contractions were dependent on synaptic transmission at NMJs, 50µM of the AChR agonist d-Tubocurarine (DTC) (Sigma) was added to the co-cultures.

#### Analysis of electrophysiological measurements

Electrophysiological measurements were analysed with custom MatLab scripts. Inward currents in voltage clamp were measured by taking the minimum value of a current trace, whereas steady state outward currents were measured by averaging over a 15ms window taken 25 ms after the voltage step. Values were corrected for baseline current offset before stimulation. Individual AP properties in current clamp were obtained using sequential injection of 100 ms current steps of increasing amplitude (10 pA increments). Only the first AP at the current threshold (first step to elicit an AP) was measured. AP waveforms were extracted using the MATLAB’s findpeaks function with minimum peak Amplitude 0 mV). Extracted parameters were: Amplitude (Max amplitude – average Vm at the end of stimulus 50 ms window excluding APs), Voltage Threshold (Voltage at the time the speed of Vm rise is above 0.15 mV/ms), Width at half heigth. Input-Output parameters were obtained using sequential injection of 500 ms current steps of increasing amplitude (50 pA increments). Location of AP were extracted using MATLAB’s findpeaks function with minimum peak Amplitude 0 mV. For analysis described in Supplementary Fig.2, firing patterns were classified as: **no AP** (No AP detected at any current injection), **single AP** (maximum 1 AP detected at any stimulation intensity), **adaptive trains of APs** (multiple APs detected at at least one stimulation intensity, but frequency decrease with increasing stimulation) and **mature repetitive AP firing** (AP frequency increases monotonically with increasing stimulation strength, without frequency adaptation). Cells with a series resistance greater than 30MΩ or a holding current lower than −100pA were rejected.

#### Statistics

Statistical analysis was performed using Prism 9 (GraphPad) and MATLAB. One-Way-ANOVA with Dunnet’s test for comparisons, Two-Way-ANOVA, unpaired, non-parametric t-tests and Mann Whitney tests were used to infer statistically significant differences between samples and groups of samples. P-values <0.05 were deemed to be statistically significant and are denoted by*. **p<0.01, ***p<0.001, ****p<0.0001. All values are represented as the mean ± SEM.

## Supplementary information

**Supplementary Figure 1.**
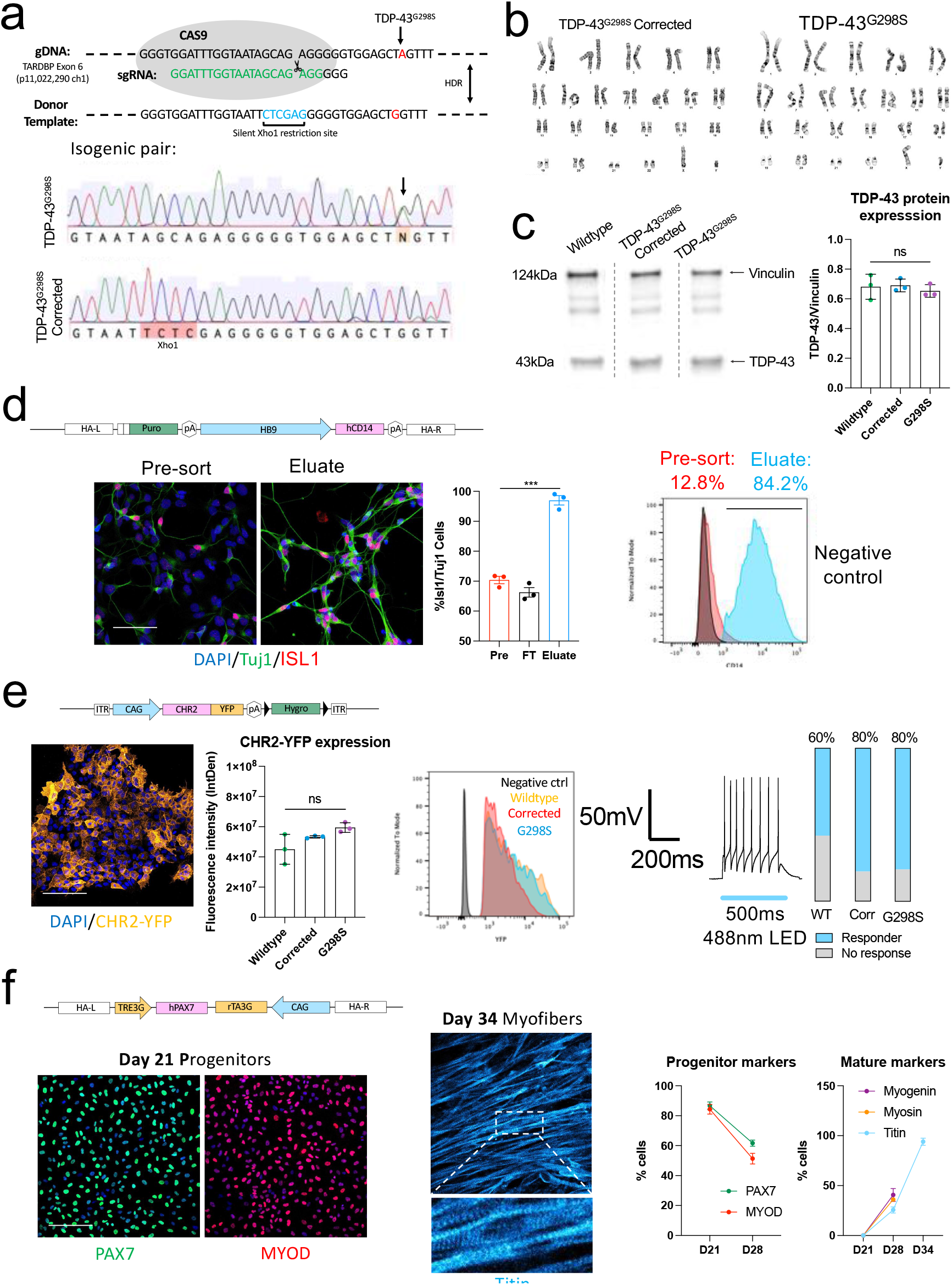
Generation of genetically engineered hiPSC lines. **A**, CRISPR-Cas9 mediated genome correction of TDP-43^G298S^ mutation. Schematic showing editing strategy and Sanger sequencing showing integration of corrected allele and silent Xho1 restriction site. **B**, G-banding showing normal karyotype following CRISPR editing **C**, Western blot showing comparable protein expression of TDP-43. D, hiPSC-lines engineered to stably express HB9::CD14 MACS sortable construct. Immunofluorescence staining and quantification for MN-specific ISL1 enrichment following MACS enrichment. Flow cytometry analysis showing MACS enrichment of HB9::CD14 MNs. Scale bar = 50μm. **E**, hiPSC-lines engineered to stably express the optogenetic construct CHR2-YFP. Flow cytometry and immunofluoresence quantification showing comparable CHR2-YFP expression across cell lines. hiPSC-MNs expressing CHR2-YFP reliably fire trains of action potentials in response to blue light (488nm) stimulation. Quantification of proportion of cells that fire APs in reponse to optogenetic stimulation. Scale bar = 50μm. **F**, hiPSC-lines engineered to stably express DOX-inducible PAX7 overexpression construct for forward programming hiPSCs into myoblasts. Immunofluorescence images of myoblast progenitors expressing PAX7 and MYOD at D21, followed by Titin expression and myoblast fusion at D34. Quantification of early and mature myogenic markers from immunofluorescence images. Scale bar = 50μm. ***p<0.001, error bars represent SEM.

**Supplementary Figure 2.**
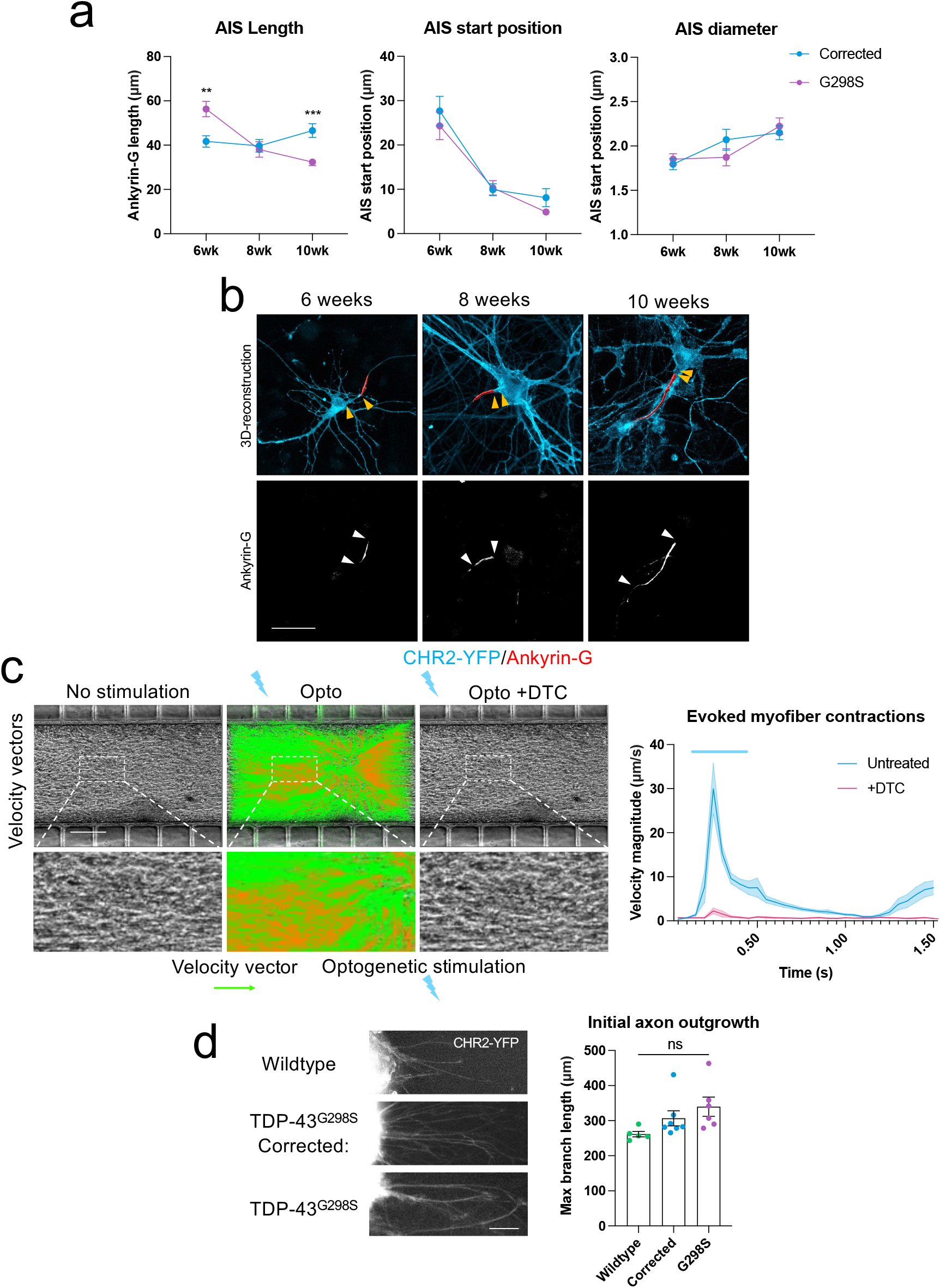
Additional AIS and neuromuscular co-culture parameters. **A**, Quantification of AIS length, start position and diameter at 6, <u>8</u> and 10 weeks maturation. **B**, Immunofluorescence images and reconstructions of the AIS based on Ankyrin-G staining at different timepoints. White arrows indicate AIS length, while yellow arrows indicate AIS start position relative to the soma. Scale bar = 50μm. **C**, PIV analysis of optogenetically evoked myofiber contractions in TDP-43^G298S^ corrected neuromuscular co-cultures before and after addition of d-Tubocurarine (DTC). Scale bar = 200μm. **D**, Quantification of D1 axon outgrowth in the neuromuscular co-cultures based on CHR2-YFP expression across genotypes. Scale bar = 50μm. Error bars represent the SEM. p-values from non-parametric, unpaired t-tests: **p<0.01, ***p<0.001, ****p<0.0001.

**Supplementary Figure 3.**
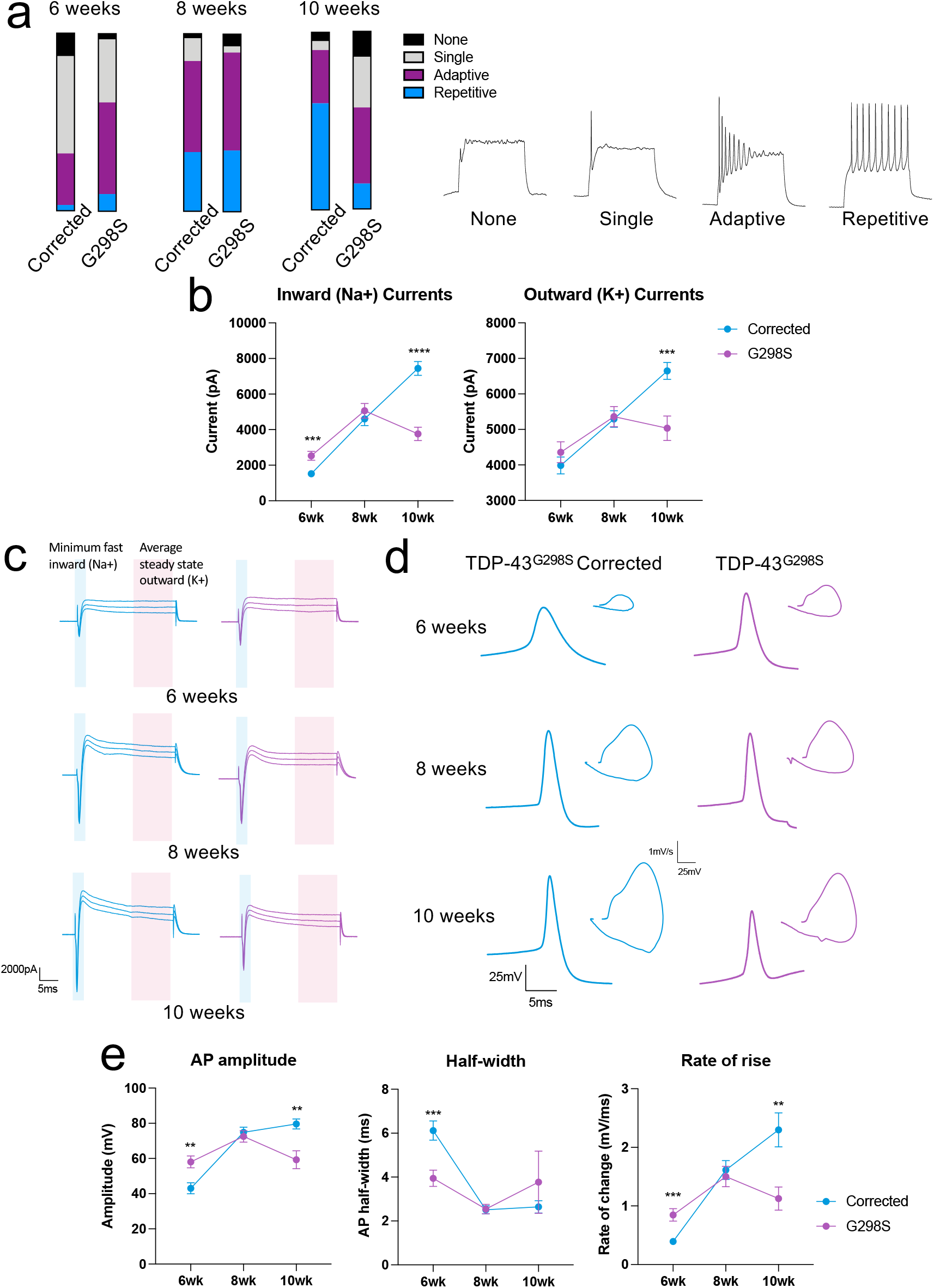
Changes in electrophysiological parameters overtime. **A**, Characterisation of firing patterns between TDP-43^G298S^ and CRISPR-corrected isogenic control lines taken at 6, 8 and 10 weeks maturation from whole cell patch clamp recordings. Firing types include: no AP, single AP, adaptive trains of APs and mature repetitive AP firing. **B**, Quantification of inward (Na+), and outward (K+) currents 6, 8 and 10 weeks maturation. **C**, Representative inward (Na+) and outward (K+) current traces overtime taken at 0,10, and 20mV pulses. **D**, Representative single AP traces and AP phase plots overtime. **E**, Quantification of AP amplitude, AP half-width and rate of change at 6, 8 and 10 weeks maturation. Error bars represent the SEM. p-values from non-parametric, unpaired t-tests: **p<0.01, ***p<0.001, ****p<0.0001.

**Supplementary Table 1.** Passive membrane properties and other electrophysiological parameters. Access resistance (RS), membrane resistance (RM), capacitance (CM), resting voltage, current threshold, voltage threshold, baseline voltage, holding current and holding voltage in wildtype, Corrected and TDP-43^G298S^ MNs at 6, 8 and 10 weeks maturation with and without optogenetic stimulation. P-values from unpaired nonparametric t-tests and one-way ANOVA tests. *p<0.05, **p<0.01, ***p<0.001.

| **6 weeks:** | **Wildtype** | **TDP-43<sup>G298S</sup>**<br>**Corrected** | **TDP-43<sup>G298S</sup>** | **TDP-43<sup>G298S</sup>**<br>**Corrected**<br>**+Opto** | **TDP-43<sup>G298S</sup>**<br>**+Opto** | **One-way**<br>**ANOVA** |
| --- | --- | --- | --- | --- | --- | --- |
| **RS** | 16.28 ±<br>1.67MΩ | 16.70 ±<br>3.84MΩ | 16.94 ±<br>1.48MΩ | 14.58 ±<br>1.54MΩ | 14.28 ±<br>2.07MΩ | p=0.87 |
| **RM** | 1063 ±<br>141MΩ | 714 ± 69MΩ | 786 ±<br>106MΩ | 1207 ±<br>314MΩ | 911 ±<br>176MΩ | p=0.09 |
| **CM** | 20.67 ±<br>2.49pF | 23.49 ±<br>2.02pF | 19.79 ±<br>3.51pF | 25.96 ±<br>2.71pF | 30.10<br>±3.60pF | p=0.13 |
| **Resting**<br>**voltage** | -35.95 ±<br>2.85mV | -38.71 ±<br>1.20mV | -38.91 ±<br>2.84mV | -35.78 ±<br>2.25mV | -34.09 ±<br>2.90mV | p=0.64 |
| **Current**<br>**Threshold** | 71.43 ±<br>10.47pA | 65.36 ±<br>6.14pA | 44.29 ±<br>3.20pA | 94.00 ±<br>10.60pA | 65.56 ±<br>9.26pA | ***<br>p=0.0007 |
| **Voltage**<br>**Threshold** | -25.07 ±<br>2.12mV | -25.11 ±<br>1.32mV | -26.55 ±<br>1.40mV | 23.47 ±<br>1.98mV | 29.40 ±<br>1.69mV | p=0.20 |
| **Holding**<br>**current** | -33.68 ±<br>4.78pA | -32.14 ±<br>5.08pA | -29.03±<br>6.81pA | -37.71 ±<br>5.59pA | 36.67 ±<br>6.29pA | P=0.77 |
| **Holding**<br>**voltage** | -59.77 ±<br>0.76mV | -60.43 ±<br>0.91mV | -60.69 ±<br>0.79mV | -60.27±<br>1.11mV | -60.90 ±<br>0.52mV | P=0.92 |

| **8 weeks:** | **TDP-43<sup>G298S</sup>**<br>**Corrected** | **TDP-43<sup>G298S</sup>** | **Unpaired T-**<br>**test** |
| --- | --- | --- | --- |
| **RS** | 10.54 ±<br>0.47MΩ | 9.80 ±<br>0.42MΩ | P=0.27 |
| **RM** | 497 ± 42MΩ | 357 ± 28MΩ | *P=0.013 |
| **CM** | 70.78 ±<br>5.73pF | 84.8 ±<br>6.10pF | P=0.10 |
| **Resting**<br>**voltage** | -51.62 ±<br>1.58mV | -43.75 ±<br>2.91mV | *P=0.01 |
| **Current**<br>**Threshold** | 77.57 ±<br>5.96pA | 89.23 ±<br>7.19pA | P=0.22 |
| **Voltage**<br>**Threshold** | -34.9 ±<br>1.40mV | -32.65 ±<br>1.37mV | P=0.27 |
| **Holding**<br>**current** | -31.41 ±<br>4.39pA | -51.07 ±<br>6.16pA | **P=0.0094 |
| **Holding**<br>**voltage** | -60.77 ±<br>0.34mV | -60.42 ±<br>0.39mV | P=0.51 |

**Supplementary Table 1. Passive membrane properties and other electrophysiological parameters.**
| **10 weeks:** | **TDP-43<sup>G298S</sup>**<br>**Corrected** | **TDP-43<sup>G298S</sup>** | **Unpaired T-**<br>**test** |
| --- | --- | --- | --- |
| **RS** | 9.06 ±<br>0.40MΩ | 11.57 ±<br>0.78MΩ | **p=0.007 |
| **RM** | 320 ± 36MΩ | 427 ± 72MΩ | p=0.20 |
| **CM** | 88.67 ±<br>7.40pF | 53.11 ±<br>6.43pF | **p=0.001 |
| **Resting**<br>**voltage** | -53.66 ±<br>2.30mV | -43.44 ±<br>3.96mV | **p=0.002 |
| **Current**<br>**Threshold** | 100.00 ±<br>9.67pA | 74.71 ±<br>9.59pA | p=0.073 |
| **Voltage**<br>**Threshold** | -31.95 ±<br>1.11mV | 29.11 ±<br>2.3mV | p=0.248 |
| **Holding**<br>**current** | -26.25 ±<br>7.36pA | -47.62 ±<br>9.28pA | p=0.08 |
| **Holding**<br>**voltage** | -60.29 ±<br>1.17mV | -60.55 ±<br>1.10mV | p=0.87 |

**Supplementary Figure 4.**
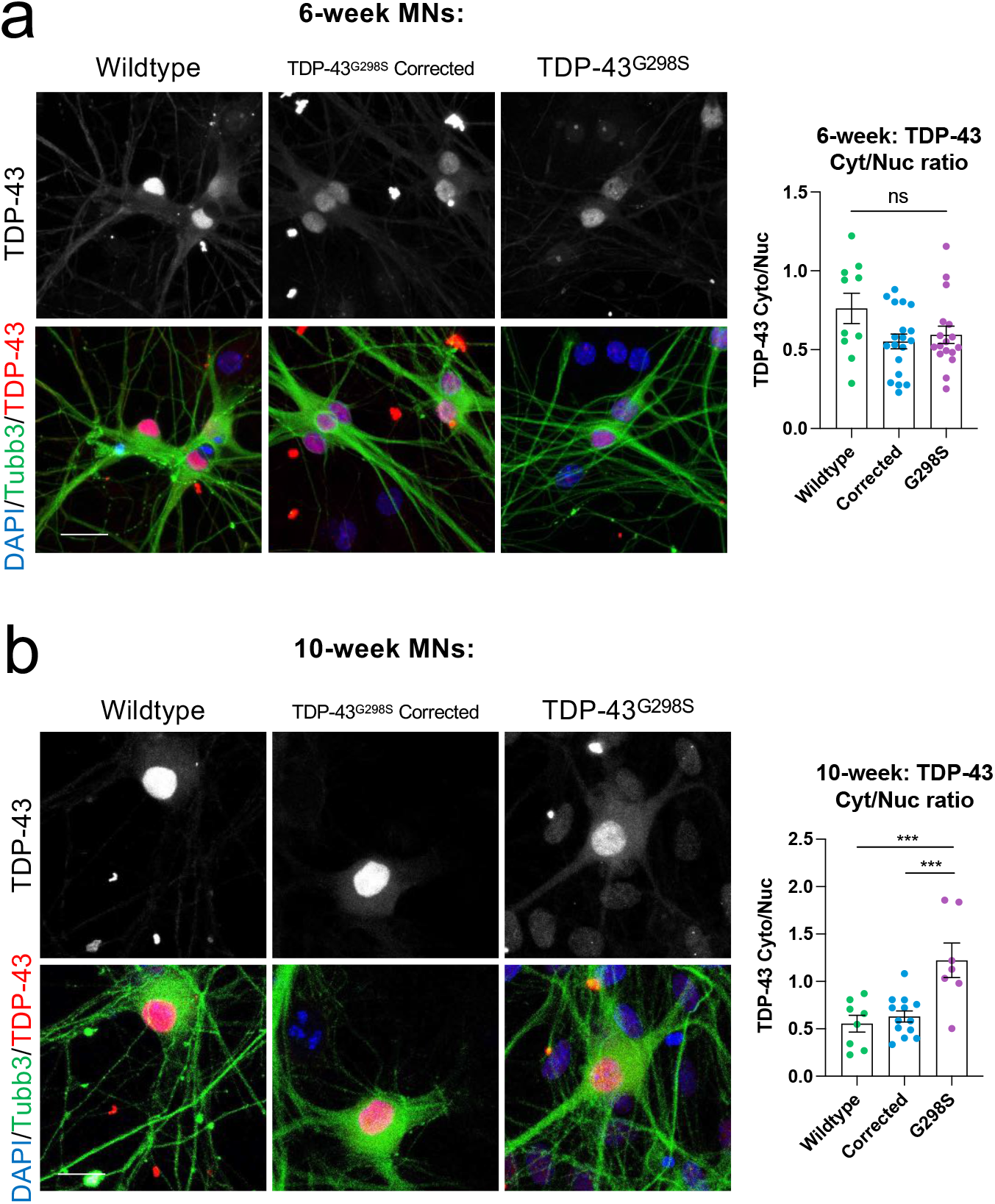
Nuclear to cytoplasmic ratio of TDP-43 in early and late MNs. **A**, Immunofluorescence images of TDP-43 (red) counterstained with Tubb3 (green) and DAPI (blue) in early (6-week) MACS enriched hiPSC-derived wildtype, corrected and TDP-43^G298S^ MNs (Scale bar = 25μm). Quantification of the ratio of cytoplasmic to nuclear fluorescence intensity in early (6-week) MNs. **B**, Immunofluorescence images of TDP-43 (red) counterstained with Tubb3 (green) and DAPI (blue) in late (10-week) MACS enriched hiPSC-derived wildtype, corrected and TDP-43^G298S^ MNs (Scale bar = 25μm). Quantification of the ratio of cytoplasmic to nuclear fluorescence intensity in late (10-week) MNs. Error bars represent the SEM. One-way-ANOVA with Dunnet’s multiple comparisons used to determine statistical significance. ***p<0.001.

**Supplementary Figure 5.**
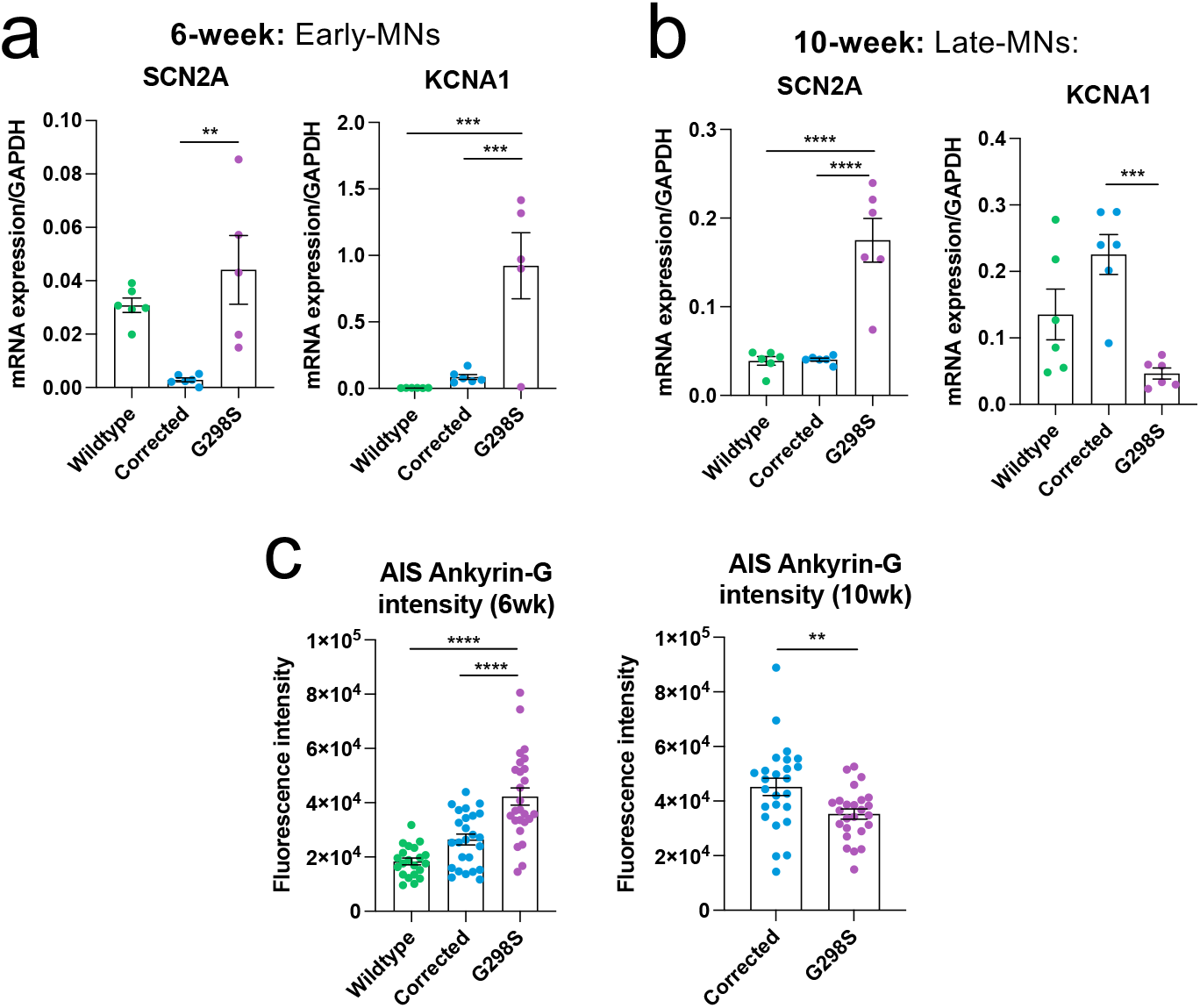
Additional AIS gene expression and Ankyrin-G protein intensity analysis. **A**, qRT-PCR analysis of SCN2A (Nav1.2), and KCNA1 (Kv1.1) expression in early (6-week motor neurons. **B**, qRT-PCR analysis of SCN2A, and KCNA1 expression in late (6-week motor neurons. **C**, Ankyrin-G staining intensity at the AIS in early and late MNs. Error bars represent the SEM. p-values from one-way ANOVA tests with Dunnet’s comparison. **p<0.01, ***p<0.001, ****p<0.0001.

## Notes

### Competing Interest Statement

The authors have declared no competing interest.

